# Synergistic interactions between confinement and macromolecular crowding spatially order transcription and translation in cell-free expression

**DOI:** 10.1101/445544

**Authors:** S. Elizabeth Norred, Rosemary M. Dabbs, Gaurav Chauhan, Patrick M. Caveney, C. Patrick Collier, Steven M. Abel, Michael L. Simpson

## Abstract

Synergistic interactions between macromolecular crowding and confinement spatially organize transcription and translation in cells. Yet, reproducing such spatial ordering in cell-free expression platforms has proven to be elusive. Here we report crowding- and confinement-driven spatial self-organization of cell-free expression that mimics expression behavior within and around the nucleoid of prokaryotes. These experiments use Ficoll-70 to approximate cellular macromolecular crowding conditions within cell-size lipid vesicles. Intriguingly, there was an abrupt change in transcriptional dynamics when crowding reached physiologically relevant levels. Imaging experiments revealed that this change in transcriptional dynamics was coincident with localization of plasmid DNA and mRNA at the vesicle wall. Computer simulations demonstrated that crowding leads to an entropically induced attraction between plasmid DNA and the wall, causing localization of DNA near the wall at sufficiently high crowding levels. The experiments demonstrate cell-like spatial organization of translation, where translational activity is controlled by chromosomally-templated positioning of mRNA. This cell-free system provides a flexible experimental platform to probe the underlying mechanisms of self-organization of membrane-less structures in cells and the spatial control of gene expression.

## INTRODUCTION

Cellular volumes are confined in a range from roughly one femtoliter^1^ to several picoliters^2^, and much of this volume (e.g. approximately 30% in *E. coli*) is occupied by proteins and other macromolecules^3–5^. The physical consequences of macromolecular crowding and cell-relevant confinement has dramatic effects on complex molecular processes, especially ones with diverse molecular components and reaction requirements like gene expression. Cell-free gene expression studies have provided a detailed (expression levels, noise, burst parameters) picture of how confinement alone ^6,7^ or crowding alone^8–10^ affect gene expression bursting (Fig. 1). Unfortunately, little is known about how synergistic interactions between confinement and crowding^11^ affect expression. It is an intriguing possibility that crowding and confinement together may have surprising effects on the complex and multi-component diffusion, binding, and reinitiation events of gene expression.

**Figure 1:**
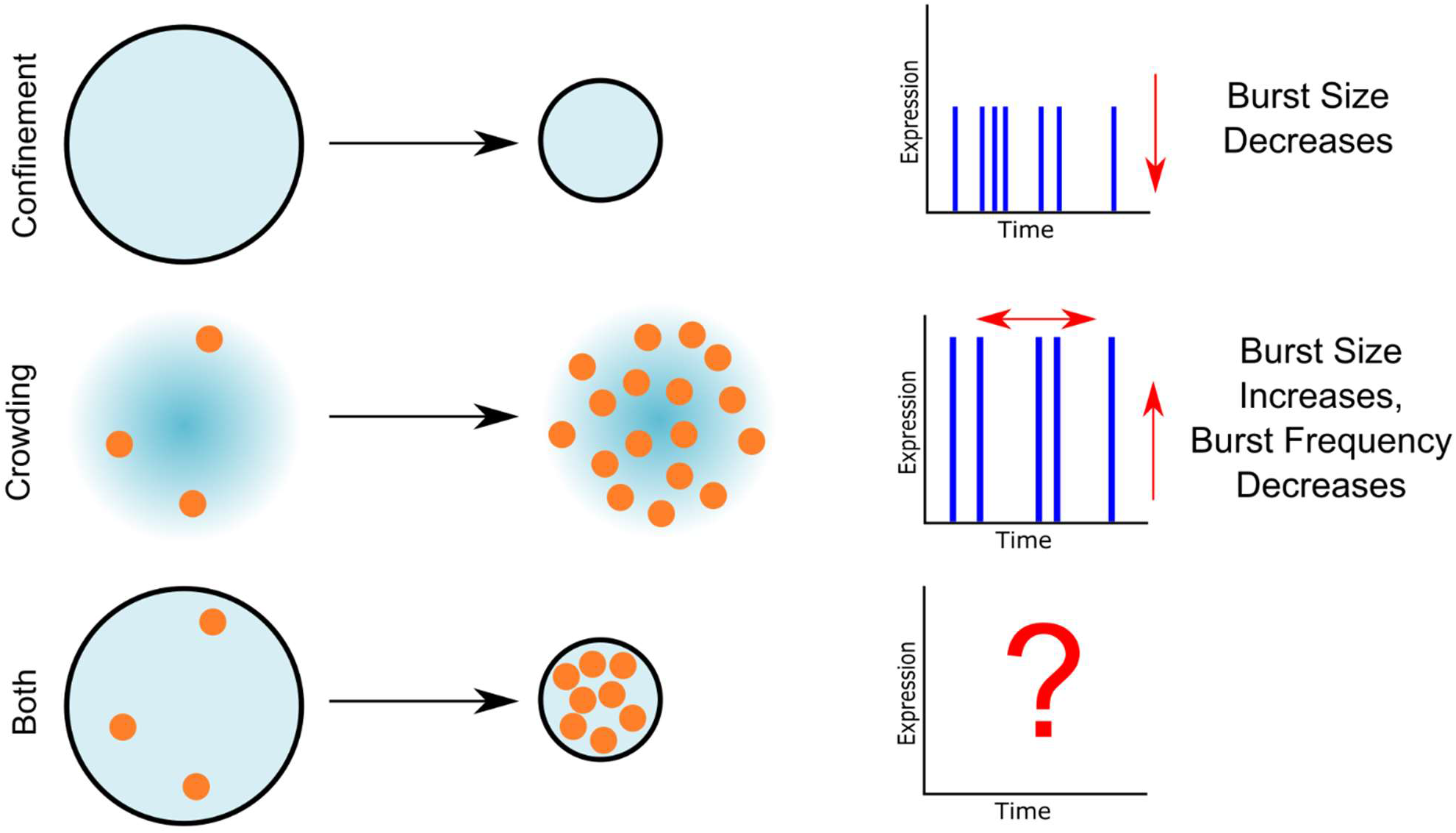
Confinement and Crowding affect gene expression bursting parameters. Gene expression bursting is sensitive to confinement (Top) and crowding (middle), but little is known about synergistic effects between crowding and cell-like confinement (bottom).

Numerous studies in confined and crowded cellular environment demonstrate how the gene expression process self-organizes into spatial subregions^12–14^. Superresolution microscopy in *E.coli* shows that transcriptional and translational components localize preferentially in different microenvironments^14^, and that transcripts often remain localized near their origin^13^. In eukaryotes, regulated phase transitions can drive spatial organization of super-enhancers that control transcriptional behavior ^15,16^. This self-organized, membrane-less structure in cells creates heterogeneous environments of crowding and confinement that control the sharing of gene expression resources and tune the patterns of expression bursting^13,14,17–19^. Cell-free systems can mimic some physical features of cells^6,9,12,20,21^, and crowding studies lacking cell-relevant confinement show some spatial organization of transcription^8,9^. Yet, more fully mimicking cell-like self-organization has been elusive.

Here we report synergistic interactions between macromolecular crowding and confinement of cell-free expression in vesicles that mimics aspects of spatial self-organization observed in prokaryotic cells. Ficoll-70 was used to approximate cellular macromolecular crowders, and crowding levels were varied from 0 to 90 mg/mL. Intriguingly, there was an abrupt change in transcriptional dynamics as crowding reached physiologically relevant levels (>40 mg/mL). Imaging experiments showed that localization of plasmid DNA and mRNA near the vesicle wall generated the change in transcriptional behavior. Computer simulations demonstrated that crowding leads to an entropically induced attraction between plasmid DNA and the wall, causing localization of DNA near the wall at sufficiently high crowding levels. At these higher crowding levels, the mRNA remained localized in the dense DNA region at the vesicle walls and was largely inaccessible for translation. These results demonstrate the spatial organization of transcription and translation in a cell-free platform that mimics the behavior within and around the nucleoid of prokaryotes, where translational activity is controlled by chromosomally-templated positioning of mRNA^13^. This work demonstrates a flexible experimental platform to understand the underlying mechanisms of selforganization of membrane-less structures in cells and the spatial control of gene expression.

## RESULTS

To understand how the combination of crowding and confinement affects gene expression, we performed cell-free protein synthesis (CFPS) reactions in vesicles crowded with Ficoll-70. Transcription and translation were tracked simultaneously using a coupled mRNA/protein reporter technique described in previous work^9,22–24^. Briefly, Spinach2^25^, an RNA aptamer which fluoresces in the green range upon hybridization with the fluorophore DFHBI-1T, was inserted downstream of a gene coding for a red fluorescent protein, mCherry^26^ (Fig. 2A). The Spinach2 fluorescence intensity was indicative of the mRNA population and transcriptional dynamics, while the mCherry fluorescence intensity was indicative of the protein population and total (transcriptional and translational) expression dynamics. Ficoll-70 at concentrations from 0-90 mg/mL was added to the Cell-free Protein Synthesis (CFPS) reactions. The concentrations of Ficoll used here mimics lower levels of physiological macromolecular crowding, which can range from 50 to 400 mg/mL^27^. Polydisperse vesicles containing the CFPS reactions were fabricated using a shearing method adapted from Nishimura *et al.* 2012^28^ (Fig. 2C). Vesicles between 14-16 μm in diameter were observed using confocal microscopy over 6 hours (Fig. 2D). Spinach2 and mCherry fluorescence were measured for individual vesicles over time (Fig. 2B, 2E). Each experiment was performed in duplicate on separate days for Ficoll-70 concentrations of 0, 10, 40, 60 and 90 mg/mL. Between 93 and 191 vesicles were analyzed per crowding condition, for a total of 694 vesicles. Transcriptional and total expression transients were extracted from individual vesicles using custom MATLAB code for image processing. The expression noise was extracted from these transients using a protocol described in previous work^6,9,12,29^ (Fig. 2E; Methods).

**Figure 2:**
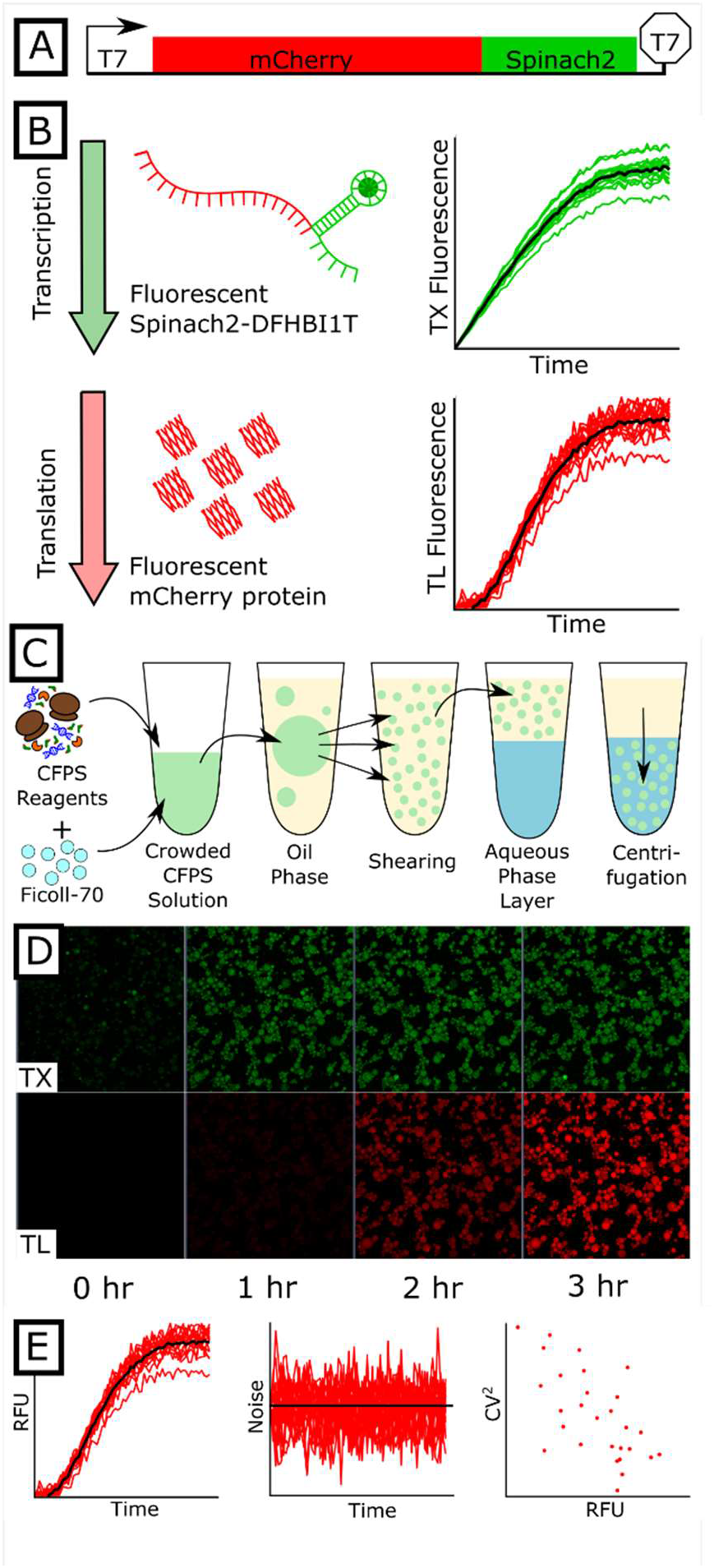
Observation of transcription and translation in cell-free reactions. A) The plasmid used for these experiments included a T7 promoter, a gene coding for mCherry, and a sequence encoding an untranslated RNA aptamer, Spinach2. B) Transcription was tracked over time by measuring the fluorescence from Spinach2-DFHBI-1T, the fluorescent hybrid of the Spinach2 aptamer and DFHBI-1T. Total expression was tracked over time by measuring the fluorescence from mCherry. C) Fabrication steps for forming vesicle microreactors. Cell-free reagents and Ficoll-70 were placed in an oil phase solution containing phospholipids, sheared into polydisperse vesicles by vortexing, layered onto a balanced aqueous phase solution, and centrifuged into the solution. D) Confocal images over time of both mRNA and protein expression in a field of polydisperse vesicles. E) Protein and mRNA expression (fluorescence of mCherry and Spinach2 in Relative Fluorescence Units (RFU)) and noise were tracked over time in individual vesicles.

In contrast to either confinement^6^ or crowding^9^ alone, the shape and timing of the transcriptional transient response varied significantly as crowding was increased in a confined environment (Fig. 3A). With confined crowding, transcription started without delay and persisted over a 100-200 minute (crowding level dependent) duration, at which point the Spinach2-DFHBI1T fluorescence reached its peak value (Fig. 3A). After the cessation of transcription, the Spinach2-DFHBI1T fluorescence decayed due to photobleaching (Fig. 3A). At the lower crowding levels (0-40 mg/mL), increased crowding decreased the transcriptional transient risetime to its peak value (Fig. 3A), with the 40 mg/mL trace reaching its peak value ~125 minutes sooner than the 0 mg/mL trace. Surprisingly, increasing the crowding level beyond 40 mg/mL reversed this trend, with the 60 and 90 mg/mL traces having risetimes similar to the 0 mg/mL transient.

**Figure 3:**
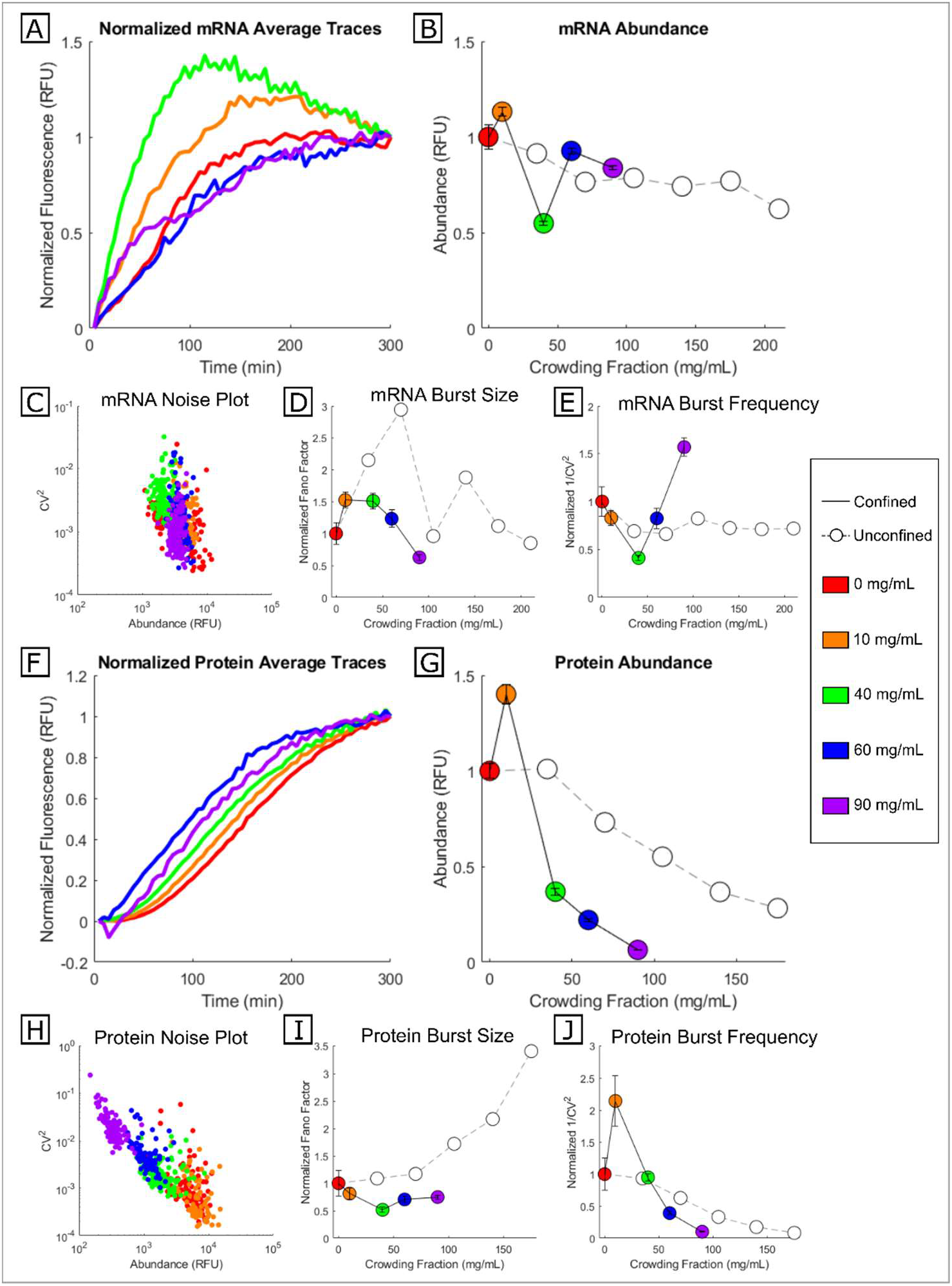
Transcription and translation in confined reaction chambers. A) Normalized average mRNA expression transient (Relative Fluorescence Units (RFU) versus crowding fraction). B) Normalized peak mRNA abundance in confined (colored dots) and unconfined (white dots) reactions. Error bars indicate standard error of the mean. C) mRNA expression noise vs. peak mRNA abundance. D) Normalized transcriptional burst size (i.e. mRNA Fano Factor) for confined and unconfined reactions. E) Normalized transcriptional burst frequency (1/CV^2^) for confined and unconfined reactions. F) Normalized average protein expression transient versus crowding fraction over time. G) Normalized peak abundance of protein in confined (colored dots) and unconfined (white dots) reactions. H) Protein expression noise (CV^2^) vs. peak protein abundance. I) Normalized total expression burst size (protein Fano Factor) for confined and unconfined reactions. J) Normalized total expression burst frequency (1/CV^2^) for confined and unconfined reactions. All unconfined batch reactions performed in a microplate reader (data taken from Norred *et al.* 2018)^9^.

A one-way ANOVA showed that confined crowding resulted in statistically significant differences in mRNA concentrations across the different crowding conditions (F(689,4)=61.47, p<0.001). In contrast to the unconfined condition^9^, the mean mRNA population was quite sensitive to crowding with confinement (Fig. 3B). Even relatively high levels of unconfined crowding (175 mg/mL) only reduced the mRNA population by about 20%, while a low level of confined crowding (40 mg/mL) reduced the mRNA population by nearly 2-fold (Fig. 3B). Surprisingly, the mRNA population did not decrease monotonically with increasing crowding fraction. Instead, a crowding level of 40 mg/mL produced the lowest mRNA population (Fig. 3B) even though this condition produced the quickest risetime.

The protein transients exhibited a delayed start in fluorescence – indicative of the maturation time of mCherry^9,26^– followed by a smooth ~250 minute rise to a peak value. In contrast to unconfined crowding^9^, mCherry maturation was significantly altered by confined crowding. The highest levels of crowding decreased maturation time by ~40 minutes (Fig. 3F) but did not otherwise significantly alter the shape of the mCherry transient (See Supplementary Information, Fig. S1).

A one-way ANOVA showed that confined crowding produced statistically significant differences in protein concentration across crowding levels (F(689,4)=526.86, p<0.001). Increased crowding reduced protein synthesis in both unconfined and confined conditions, but cell-like confinement significantly amplified the effects of crowding (Fig. 3G). Compared to no crowding, confined crowding of 90 mg/mL reduced protein production by more than an order of magnitude. In contrast, a similar decrease in protein population required an unconfined crowding level exceeding 175 mg/mL (Fig. 3G). Consistent with other reports^10^, there was a statistically significant 1.4 fold increase in protein synthesis with a low level (10mg/mL) of confined crowding (Fig. 3G).

The details of expression behavior were investigated by examining the noise behavior of the two reporters using the relationships:

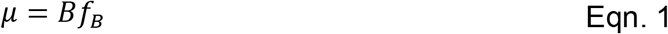

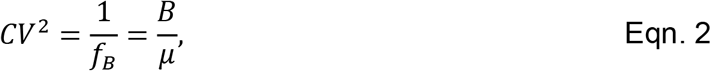

where *μ* is the mean population of the reporter (i.e. mCherry or Spinach2-DFHBI1T); *CV*^2^ is the square of the coefficient of variation (variance of reporter population/ *μ*^2^); and B and fB are parameters that describe the expression pattern. In the 2-state model of expression bursting from an individual gene, B is the burst size (average number of molecules created per burst) and f_B_ is the burst frequency (number of bursts per unit time)^9,30–37^. With multiple copies of plasmids in each vesicle, the burst frequency may be thought of as the number of statistically independent expression centers, and the burst size as the intensity of expression within the centers^12^. There is evidence of expression patterns indicative of these distinct expression centers even without crowding^12^, but with crowding these centers (at least at the transcriptional level) are visible using optical microscopy^8,9^. The transcriptional and total expression burst sizes (*B* = *μCV*^2^ (also known as the Fano factor); Fig. 3D and 3I) and frequencies 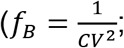 Fig. 3E and 3J) were calculated using the measured values of *μ* and *CV*^2^ for each reporter in every vesicle (Fig. 3C and H).

There was a drastic shift in transcriptional burst behavior between the lower and higher crowding levels. The change in transcriptional burst size (B_TX_) across all crowding levels with confinement was statistically significant as determined by one-way ANOVA (F(689,4)=11.48, p<0.001). B_TX_ increased by ~1.5 fold in response to low levels of confined crowding (10, 40 mg/mL; Fig. 3D), but decreased at higher (≥ 60 mg/mL) crowding levels (Fig. 3D). The abundance of transcriptional expression centers (f_BTX_) had a statistically significant change across all crowding levels as determined by one-way ANOVA (F(689,4)=50.38, p<0.001). As the crowding level increased from 0 to 40 mg/mL, f_BTX_ decreased by ~5 fold. Yet at higher levels of crowding (≥ 60 mg/mL), f_BTX_ exceeded the value measured with no crowding (Fig. 3E).

Similar to unconfined crowding, there was little change in the total expression burst size (B_T_; protein Fano factor) for the crowding levels measured here (Fig. 3I). Although this change in the protein Fano factor was marginally statistically significant as determined by one-way ANOVA (F(689,4)=2.77, p=0.0263), only the two groups that produced the highest and lowest protein Fano factor (0 mg/mL and 40 mg/mL) were significantly different from each other. This is similar to unconfined reactions, where protein Fano factors remained relatively unchanged for crowding <140 mg/mL, although higher levels of unconfined crowding did result in large increases (~4-fold) in the protein Fano factor^9^. In contrast, the abundance of total translational expression centers (f_BT_) was quite sensitive to the crowding level (Fig. 3J) and varied by more than 30-fold from its peak value at low crowding (10 mg/mL) to its lowest value at high crowding (90 mg/mL). This change was statistically significant as determined by one-way ANOVA (F(689,4)=154.38, p<0.001). As found for protein concentration, confinement amplified the crowding induced decrease in f_BT_ (Fig. 3J).

Intriguingly, the results here show a decoupling between transcriptional and translational expression centers. While all translational expression centers must be initiated by a transcriptional center, many transcriptional centers were not translationally active. For example, the spike in protein population with 10 mg/mL confined crowding (Fig. 3G) was much larger than the associated mRNA concentration increase (Fig. 3B) because this low level of crowding increased the number of transcriptional expression centers (as indicated by f_BTX_) that were translationally active (as indicated by f_BT_; Fig. 3J). Yet, as the crowding level was increased, transcriptional expression centers became more elusive for the translational machinery. The abundance of transcriptional expression centers reached a peak with 90 mg/mL of confined crowding, but nearly none of these centers were translationally active (Fig. 3J).

Previous reports demonstrated that crowding without cell-relevant confinement affected translational activity by creating an inhomogeneous spatial distribution of mRNA^8,9^, and with increased crowding, much of the mRNA became inaccessible for translation. An especially intriguing feature of expression with confined crowding was the abrupt change in transcriptional behavior as crowding increased from 40 to 60 mg/mL (Fig. 3B and 3E), implying a pronounced shift in the mRNA spatial organization. To examine the evolution of mRNA spatial organization with increased crowding, representative vesicles of the same approximate size were compared visually using confocal microscopy. Fig. 4A shows fluorescence cross-sections of vesicles of ~15um diameter at the endpoint of a 300-minute reaction. With no crowding, mRNA was visible in distinct spots of relatively uniform intensity and spatial distribution (Fig. 4A). Low levels of crowding (10 mg/mL) had little discernable effect on the spatial distribution of the mRNA, but as we previously reported^9^ did lead to the emergence of a few hot spots of higher local mRNA concentration (Fig. 4A). At a crowding level of 40 mg/mL these hot spots preferentially appeared near the walls of the vesicles, and were almost exclusively found at the walls with crowding of 60 mg/mL (Fig. 4A). This localization of mRNA at the vesicle walls was not seen in larger (~90 μm diameter) cell-free reaction chambers^8^, indicating that the synergistic effects of confinement and crowding^11^ emerge at cell-relevant confinement volumes.

**Figure 4:**
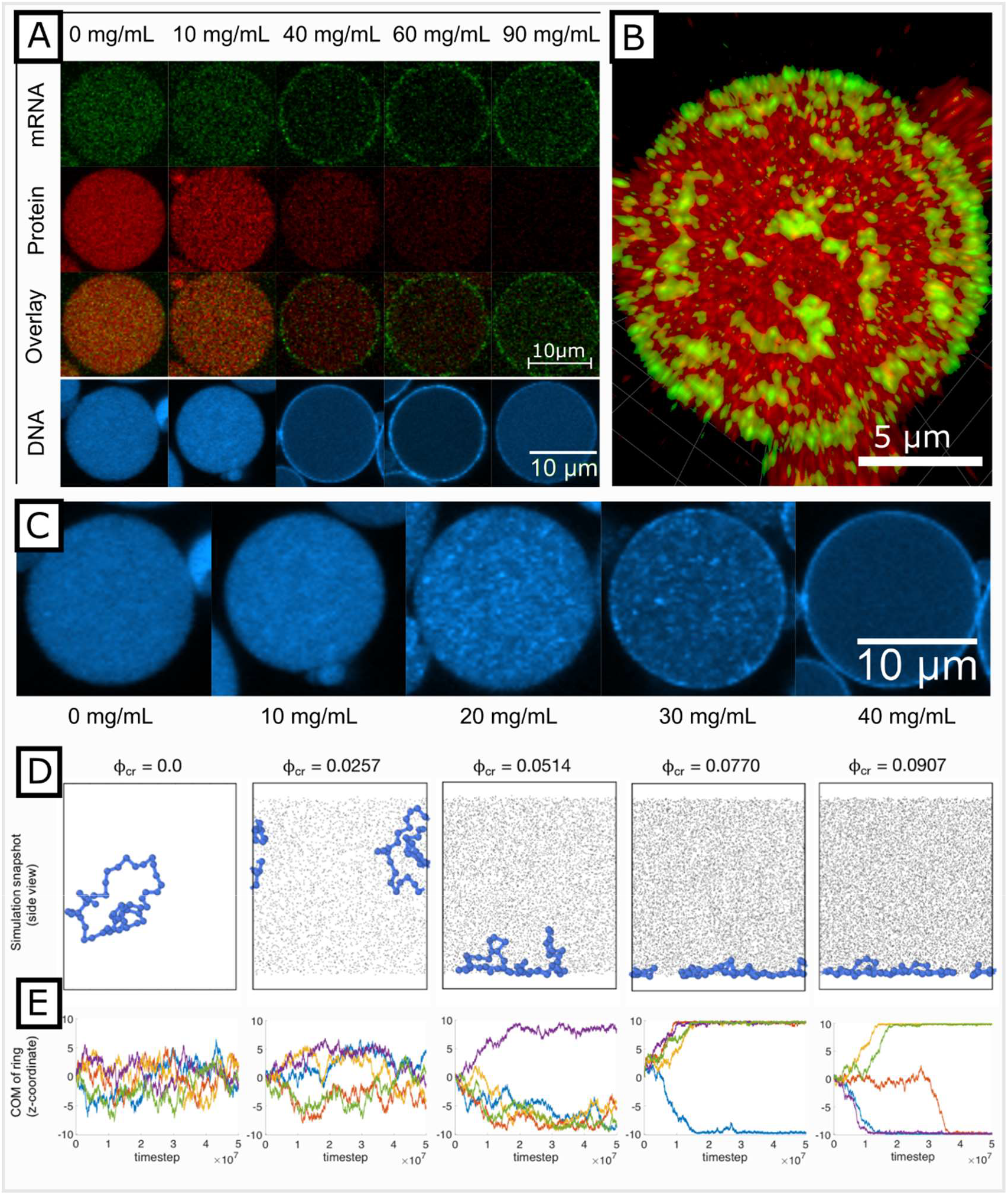
Spatial distribution of protein, mRNA, and DNA in vesicles. A) Representative vesicles demonstrating spatial distribution of mRNA and protein. Overlay shows combination of mRNA and protein signals in the same vesicle. Bottom row shows representative vesicles from separate experiments where DNA was fluorescently labelled with Pico488 dye (false colored to cyan). B) Three dimensional z-stack reconstruction of an individual vesicle. C) Distribution of DNA demonstrated by fluorescent labelling with Pico488 (false colored to cyan). D) Representative snapshots from equilibrated simulation trajectories at five different crowding fractions (DNA in blue, crowders in grey). The system is viewed from the side and is confined at the top and bottom. E) The z-component of the center of mass of the polymer, with walls confining the system at z = +10 and z = −10 in reduced units. Results from five independent simulations are shown for each crowding fraction. The polymer starts from z = 0 in each simulation.

There was no evidence of mRNA diffusion to the wall after synthesis at other locations. Instead, time-lapse images (Fig. S2) indicate that with crowded confinement, transcription occurred at the wall, and the mRNA remained localized near the site of transcription throughout the experiment. A 3D reconstruction of a crowded vesicle, created from z-stack confocal imaging, showed that mRNA were synthesized in hot spots around the periphery of the vesicles (Fig 4B). Since imaging experiments in prokaryotes show that mRNA often exhibit limited dispersion from their site of transcription^13^, we hypothesized that confined crowding led to localization of the plasmid DNA near the vesicle wall. To test this hypothesis, we prepared the vesicles with a DNA dye, Pico488, and examined fluorescence cross-sections of ~15 μm diameter vesicles using a confocal microscope. These measurements showed DNA was spatially organized in the same patterns as the mRNA. Without crowders, there was a sparse distribution of DNA throughout the interior of the vesicle. As the crowding level increased, the DNA appeared in localized hot spots distributed throughout the interior of the vesicle. At a crowding level of 40 mg/mL and higher, DNA localized near the vesicle wall (Fig 4A, 4C).

We further explored this phenomenon using Brownian Dynamics computer simulations. We utilized a coarse-grained model of a DNA plasmid in a crowded and confined environment: The DNA plasmid was modeled as a flexible ring polymer, the volume fraction of crowders (ϕ_cr_) was varied by changing the number of crowding particles, and all components interacted via short-ranged repulsive interactions. The system was confined by repulsive walls in one dimension (z) and had periodic boundary conditions in the other dimensions. A full description of the model is provided in the SI. Figure 4D shows the behavior of the polymer, initially located in the interior of the system, as a function of the volume fraction of crowders. At low crowding fractions, the ring polymer adopts coiled configurations in the bulk of the system. However, at higher crowding fractions, it localizes to the wall. At ϕ_cr_ = 0.077 and higher, the polymer nearly completely flattens against the wall even though there are no specific attractive interactions between the two. The effective attraction is a consequence of depletion interactions resulting from the presence of crowding particles^38,39^. The crowding-induced localization observed in simulations is consistent with the experimental observations (Fig. 4C) in which DNA plasmids are found near vesicle walls at high crowding levels.

The cell-free results reported here are strikingly similar to expression behavior in prokaryotic cells where mRNA localization determines translational efficiency^13^. Superresolution microscopy of *E. coli* shows that high-rate transcription preferentially occurs at the periphery of the nucleoid^17^, and this mRNA population is efficiently translated as it resides at the boundary with ribosome-rich regions of the cell^14^. Conversely, lower rate transcription occurs throughout the ribosome-poor^14^ nucleoid, and the resulting relatively immobile mRNA populations are inefficiently translated^13^. At low levels of confined crowding in cell-free reactions, the mRNA is expressed in distinct translationally-active regions. At higher levels of crowding, the DNA is localized and compacted near the vesicle wall, and the resulting mRNA remains localized in this dense DNA region that appears to be largely inaccessible for translation. By generating a spatial organization of transcription and translation that mimics key aspects of prokaryotic cell membrane-less structure, these cell-free experiments provide a flexible experimental platform to probe the underlying mechanisms of cellular self-organization.

While Ficoll 70 provides a reasonable approximation of cellular crowding, there are important differences to note. An *E. coli* cell is approximately 50-70% water, and the dry weight is ~55% protein, ~20% RNA, ~10% Lipids, and ~15% of other molecules^40^. As much of the protein and RNA is ribosomal^41^, ribosomes (~20 nm in diameter in prokaryotes) are a significant contributor to cellular macromolecular crowding. Most of the non-ribosomal protein has radii in the 3-6 nm range with a globular configuration^42^. Ficoll 70 (stokes radius of ~5 nm) and the PURE expression media used here accurately approximate the distribution of globular protein and ribosome crowders in cells. However, extended structures like cytoskeletal filaments or elongated proteins, and structures like the bacterial nucleoid or eukaryotic organelles are not well approximated in these experiments. In contrast to these cell-free experiments, in cells these extended structures may affect expression bursting by allowing for facilitated transport or by creating inhomogeneous crowding^14^. Finally, the results presented here should be applied carefully concerning eukaryotic expression. First, eukaryotic transcriptional burst dynamics are highly sensitive to promoter structure (e.g. TATA boxes) or nucleosome occupancy patterns ^43^ not present in these cell-free experiments. Furthermore, eukaryotic cells completely decouple transcription and translation, and includes addition steps (e.g. mRNA export from the nucleus) that may affect expression noise^44^.

The history of cell-based synthetic biology is one of gene circuit design using specific molecular mechanisms (e.g. promoter/transcription factor interactions) and principles borrowed from electronic circuit design^45^. Much of cell-free synthetic biology has followed a similar path^46–48^. However, there is a growing realization that the manipulation of the expression environment – from the composition of the expression reaction media^49^ to the physical (confinement and crowding) arrangements – provide another dimension to cell-free synthetic biology. One big advantage of cell-free platforms is that they provide the ability to intricately vary spatial arrangements^50^ and are especially well-suited for spatial synthetic biology as a strategy to achieve specific functionality. However, the results here – which show the cell-free spatial organization of expression much like that seen in prokaryotes – suggest that the most immediate application of these experimental systems is to understand the underlying mechanisms of self-organization of collective behavior in cells.

## METHODS

In order to simultaneously track transcription and translation outputs, a plasmid vector coding for mCherry and a downstream fluorescent mRNA aptamer, Spinach2, was expressed^23,25^. The plasmid pRSET-b-mCherry-Spinach2 transcribes from a T7 polymerase promoter to create a transcript with a translated region coding for mCherry, followed by an untranslated aptamer tag which fluoresces after folding and binding with the fluorophore DFHBI-1T^51^ ((Z)-4-(3,5-difluoro-4-hydroxybenzylidene)-2-methyl-1-(2,2,2-trifluoroethyl)-1H-imidazol-5(4 H)-one, Lucerna, Inc). A commercial cell-free protein synthesis kit (PURExpress, NEB) was used to express the plasmid in the presence of DFHBI-1T and a crowding molecule, Ficoll-70 (Sigma-Aldrich).

Vesicles containing the cell-free expression system and added components (the “Inner Solution” were prepared by a shearing method adapted from Nishimura *et al.* 2012. In summary, vesicles are prepared by assembling the cell-free reaction mixture, plasmid, DFHBI-1T, sucrose (to aid with visualizing vesicles), and Ficoll-70. Concentrated Ficoll-70 was added at a final concentration from 0-90 mg/mL. The Inner Solution is vortexed in a paraffin oil solution containing phospholipids (POPC, Avanti Polar Lipids) to create a polydisperse population of water-in-oil droplets. This paraffin oil mixture with droplets is layered on to an aqueous “Outer Solution” and then centrifuged for 20 minutes at 4C at ~14k g. The Outer Solution is balanced with the inner solution, containing small molecules found in the PURE system reactions^52,53^ (See SI for list of reactants). Vesicles are collected by pipetting and are prepared for microscopy by placing ~10 μL of vesicles in Outer Solution between two glass coverslips separated by a ~2 mm PDMS spacer. Most vesicle diameters range from 5-30 um.

Vesicles were observed while resting on a coverslip, using a (Zeiss LSM 710 Axio Observer) confocal microscope to image every 5 minutes for 6 hours. A 20x objective (Zeiss Plan Apochromat 20x/0.8 M27) was used for the timescale data, followed by a 63x objective (Zeiss Plan Apochromat 63x/1.40 Oil DIC M27) for a more detailed image of fluorescence distribution at the end of the experiment. The Spinach2-DFHBI-1T signal was measured using a 561nm laser from 488/536 nm Ex/Em. The mCherry was measured using a 561 nm laser from 561/637 nm Ex/Em. Brightfield images were also acquired contemporaneously. For each timepoint, the vesicles were imaged using a z-stack capture, using ~14 images per slice at 2 um increments. The images were analyzed using ImageJ and custom MATLAB code to detect vesicle size and location and to acquire intensity values for Spinach2-DFHBI-1T and mCherry from individual vesicles over time.

To determine spatial DNA distribution in the vesicles, vesicles were prepared as normal with the 0.25 μL of the 200x fluorophore Pico488 (Lumiprobe) in the Inner Solution, instead of DFHBI-1T. These vesicles were imaged using confocal microscopy, using a 63x objective and 561 nm laser at 488/536 Ex/Em. Ficoll-70 was added at a final concentration at 0, 10, 20, 30, 40, 60, and 90 mg/mL. A control reaction containing no DNA was also performed. Z-stack renderings and cross-sections in the middle of vesicles were used to characterize DNA distribution within vesicles.

Two experiments were performed for each Ficoll-70 crowder concentration of 0, 10, 40, 60, and 90 mg/mL. For an individual vesicle in each experiment, a 6-hour mRNA and protein expression trace was extracted for noise analysis, as described in previous work^6,9,12,30^. Built-in functions in ImageJ and custom MATLAB code were used to identify the boundaries of vesicles in a brightfield view, select regions of interest around each vesicle, and extract fluorescence information from each ROI. Fluorescence for individual vesicles was tracked over time for detected vesicles between a diameter of 14-16um.

Briefly, for each population of zeroed expression traces in a single experiment, an average trace, or “general trend” was calculated for all vesicles, and then was subtracted from each individual vesicle’s expression trace. This was done for both reporters, revealing the “noise signals”, or the stochastic variation in mRNA or protein reporter at each timepoint. The coefficient of variation squared (CV^2^) was used to quantify the noise magnitude in the molecular populations of mRNA or protein. The coefficient of variation squared is defined in Eqn. 2. The steady state fluorescence level was defined as the maximum fluorescence level attained per fluorescence trace, instead of the endpoint of the trace. This was due to trace decay caused by photobleaching, causing the final fluorescence value not to be descriptive of the total molecular populations produced. Since the mRNA traces reached their final value prior to the protein traces, noise traces were only calculated based on the first 150 minutes of the mRNA reactions. However, protein noise traces, which are derived from traces that generally express continuously over the entire experiment, were calculated over 300 minutes. CV^2^ is plotted against these maximum values, which is useful for describing changes in the bursting patterns between experimental conditions.

## ACKNOWLEDGEMENTS

This research was conducted as part of the Interface Directed Assembly Theme at the Center for Nanophase Materials Sciences, which is a DOE Office of Science User Facility. SEN and PMC acknowledge Graduate Fellowships from the Bredesen Center for Interdisciplinary Research and Graduate Education, University of Tennessee, Knoxville. The authors acknowledge Drs. Roy Dar, Brandon Razooky, Maike Hansen, Leor Weinberger, Scott Retterer, and members of Dr. Mitch Doktycz’s laboratory for their helpful feedback and fruitful discussions.

